# Effects of speaker and listener sex on auditory attention decoding performance

**DOI:** 10.1101/2025.05.22.655324

**Authors:** Iris Van de Ryck, Nicolas Heintz, Iustina Rotaru, Debora Fieberg, Alexander Bertrand, Tom Francart

## Abstract

**Objectives:** Auditory Attention Decoding (AAD) is a technique utilizing brain signals to decode on which sound the listener focuses the attention. In most current studies, the effect of type of speech materials used and sex of the listener is not considered. We investigated the effect on AAD performance of factors related to the speaker (such as the sex of the speaker, background noise level, and same versus mixed-sex conditions) and the listener (sex of the listener).

**Design:** Forty-two young adults with normal hearing participated in the study. They listened to 2 competing speakers and were instructed to attend to one speaker and ignore the other speaker, whilst electroencephalography (EEG) and electrooculography (EOG) were measured. Background noise was introduced in half of the conditions. AAD performance was compared across eight experimental conditions.

**Results:** A significant main effect of speaker sex was found: A male target and/or male masker speaker resulted in higher AAD performance compared to a female speaker with a higher fundamental frequency. These effects were found to be small and therefore likely clinically irrelevant.

**Conclusion:** While no substantial effects were found on the factors investigated in this study, including diverse and realistic training scenarios remains a valuable approach to prevent potential influences from other factors.

## 1. Introduction

**Auditory attention decoding (AAD**) refers to the problem of detecting the attended speaker from measured brain signals, which could potentially be leveraged by a hearing aid’s noise reduction algorithm to enhance the attended speaker. AAD is addressed through techniques involving brain signal decoding, particularly using methods like electroencephalography (EEG). More specifically, the envelope of speech can be reconstructed from neural recordings because cortical activity phase-locks to the envelope of an attended speech source, such that the actual speech envelope can be correlated to the one derived/reconstructed from neural recordings (O’Sullivan et al., 2015). Within a multiple-speaker scenario, the speech envelope that exhibits the strongest correlation with the neural-reconstructed envelope is then selected as the attended speaker. One of the applications of AAD are neuro-steered hearing aids (e.g., Biesmans et al., 2015; Mirkovic et al., 2016; Van Eyndhoven et al., 2017).

To date, many AAD algorithms have been developed (e.g., Das et al., 2016, 2017; Geirnaert et al., 2019, 2020; Han et al., 2019; Rotaru et al., 2024; Tanveer et al., 2024; Wang et al., 2021; Wong et al., 2018; Xu, Bai, Zhao, Zheng, et al., 2022; Xu, Bai, Zhao, Hu, et al., 2022), some of them also being combined with noise suppression systems (Biesmans et al., 2017; Das et al., 2016, 2017; Van Eyndhoven et al., 2017).

In the literature, we learn that a range of **factors influence AAD performance,** but many of these factors remain unclear or unexplored. To date, factors that have been investigated are head-related filtering and ear-specific decoding bias (Das et al., 2016), distractors (Holtze et al., 2021), level of background noise (Aroudi et al., 2019; Das et al., 2018), speaker position/separation (Das et al., 2018), reverberation (Aroudi et al., 2019), interfering speakers (Aroudi et al., 2019) and speech intelligibility (Das et al., 2018). We briefly review these below.

In the current AAD literature the choice of stimuli and selection of participants has often been rather arbitrary. It has not been systematically investigated what the effect is of factors such as the **sex of the target/masker speaker, same/mixed-sex combinations of speakers and the sex of the listener** on AAD performance and **subjective experience**. This is problematic because the AAD algorithm must demonstrate accurate performance across various real-life listening situations to be able to implement the algorithm in future hearing aids.

Das et al. (2016) highlighted the importance of **head-related filtering** and **ear-specific** decoding bias, with the former significantly improving performance. Other literature looked into the effect of **distractors.** Holtze et al. (2021) found that after presenting a distractor (by embedding the name of the person in the ignored speech stream), the speech envelope tracking increased for both the attended and unattended speech. Other factors that have already been explored, include the **level of background noise** and **speaker position** (angular separation), both of which have been demonstrated to influence attention decoding accuracy (Das et al., 2018). Das et al. (2018) demonstrated that surprisingly, attention decoding performance improved when moderate background noise (-4,1 dB SNR, -1,1 dB SNR) was introduced, in contrast to no noise (silence + 20dB SNR). However, as the noise level increased to -7,1 dB SNR, decoding performance notably decreased. They hypothesize that adding a small amount of background noise increases listening effort, leading to larger brain responses and therefore improved AAD performance. On the other hand, Aroudi et al. (2019) found that **background noise** and **reverberation** did not significantly impact EEG-based auditory attention decoding, but that the presence of an additional **interfering speaker** could notably reduce decoding performance. Aroudi et al. also stated that we must take careful consideration of acoustic components such as reverberation, background noise and interfering speaker, because they are crucial for optimizing the decoding process in challenging acoustic environments. In this study, we investigated the influence of adding a small amount of background noise compared to conditions without background noise on AAD performance in normal-hearing young subjects. Extrapolating the results from (Das et al., 2018), we hypothesized that AAD performance is better in conditions with a background noise of 4 dB SNR compared to conditions without background noise, due to increased listening effort. Moreover, adding a small amount of background noise closely matches real-life listening situations.

Additionally, AAD performance can improve with increasing **speaker separation** (10°, 60°, 180°), demonstrating the benefit of spatial release from masking (Das et al., 2018). This effect of speaker separation on decoding accuracy was amplified when the level of background noise increased. The degrading effect of the noise was stronger when competing talkers were located closer to each other. Furthermore, Ericson et al. (2004) also found that behaviourally spatial separation improves the intelligibility of all the talkers in the stimulus. Moreover, Das et al. (2018) found a significant correlation between **speech intelligibility** and attention decoding performance across different acoustic conditions. (Das et al., 2018).

As stated before, many factors that could influence AAD performance remain unexplored. The effects of the **sex** of both the **speaker** and **listener** have not been explored specifically in AAD protocols. In literature, we find that sex differences can be found in the voice and speech of men and women. There are **anatomical differences** (Titze, 1989; Titze & Martin, 1998), differences in **acoustical** perception of sex in speech (F0, timbre) (J. Hillenbrand et al., 1995;J. M.Hillenbrand & Michael, 2009; Pernet & Belin, 2012), and **sociolinguistic differences** (age, prosody, intonation) (Coates, 2004; Leung et al., 2018) between males and females. Research has also been conducted about differences in how the **brain** responds (neural level) to speech of male versus female speakers (Ahadi et al., 2014; Durrant et al., 1990; Jerger & Johnson, 1988; Krizman et al., 2012; Van Canneyt et al., 2021; Weston et al., 2015; Zakaria et al., 2019), about the **social/behavioural level** (Hazan & Markham, 2004; Kiliç & Ögüt, 2004; Zekveld et al., 2014), and about **speech understanding/speech intelligibility** (Ellis et al., 1996; Kwon, 2010; Yoho et al., 2019). **Nevertheless, specifically the effect of speaker’s and listener’s sex on AAD performance remains underexplored.**

In the next two paragraphs we zoom in on the effects of speaker’s and listener’s sex on the behavioural and neural level.

Research indicates that talker and listener sex influence **behavioural** results such as **speech intelligibility**, though findings vary. A study by Ho-Beom Kwon (2010) suggested that talker sex can influence speech intelligibility, potentially due to psychological or acoustic differences between male and female speech, but there was no consensus on which sex produces the highest scores. Hazan and Markham (2004) investigated the influence of both the speaker’s and listener’s sex on speech intelligibility. Different methods for measuring intelligibility (percentage of words correct versus subjective rating scales) and data collection (laboratory versus crowdsourcing) were compared. The study showed that the speech of **female** speakers was generally more intelligible, but that there was considerable variability among speakers. Furthermore, an interaction between the sex of the speaker and listener was observed, depending on the method used. Crowdsourcing led to less bias in subjective ratings. The study emphasized the importance of considering these factors when assessing speech intelligibility. A study by Yoho et al. (2019) also explored the influence of both talker and listener sex on speech intelligibility. The study showed that, regardless of methodology, the spoken productions of **female** talkers were more intelligible than the spoken productions of male talkers. However considerable variability across talkers was observed. Yoho et al. underscored the need for researchers to consider talker and listener sex when assessing speech intelligibility and when comparing results across studies that have used different demographics and methodologies. Additionally, Ellis et al. (1996) found that **women** may be perceived as more understandable by male listeners, while men may be perceived as more understandable by female listeners. However, objectively, there was no significant difference between male and female subjects’ magnitude-estimation scaling responses. **These studies suggest that talker and listener sex can influence speech intelligibility, but there is no clear consensus on which sex induces the highest scores.**

**Listener’s** and talker’s **sex** differences have also been found on the **neural level**. Auditory brainstem responses (ABR) have been shown to be influenced by sex. There are gender-specific normative data for ABR in clinical applications (Hall, 1992). Also other studies confirm that neural encoding of the acoustic cues of speech stimuli is different at a subcortical level between male and female **subjects**. (Ahadi et al., 2014; Durrant et al., 1990; Jerger & Johnson, 1988; Krizman et al., 2012; Zakaria et al., 2019). Next to listener’s sex, also **talker’s sex** differences have been found on the neural level. Weston et al. (2015) conducted a study with MEG, showing more activation of the non-primary auditory cortex for the female timbre compared to the male timbre, primarily due to the increased temporal complexity of female voices. This increased activation associated with female voices was not solely due to fundamental frequency (F0), nor due to psychological attribution of female sex nor affected by listener sex. That study of Weston et al. (2015) investigated the perception of voice sex and showed that male and female voices are represented as distinct auditory objects in the brain. The mechanism underlying sex discrimination relies on sex-dependent activation cues in the non-primary auditory cortex. A study by Van Canneyt et al. (2021), using EEG, showed that the stories that were narrated by a male speaker (with a low and steady **fundamental frequency (F0)**) evoked stronger f0-tracking compared to female-narrated stories (with high and variable F0), for which many responses were not significant. These studies suggest that sex-related voice characteristics influence brain signals. It is not known yet whether the sex of the **speaker** affects AAD performance in a single or multiple speaker scenario. **Based on the literature, we expect to find a difference in performance between male and female speakers but not between listeners.**

In standard AAD paradigms, a minimum of 2 speakers are presented, leading to **same-sex** or **mixed-sex** situations. This has also not yet been investigated for AAD performance. For speech understanding in these situations, both **energetic** (related to acoustic energy) and **informational masking** (related to cognitive processing) play an important role (Francart et al., 2011). The effect of masking is dependent on the properties of the speech and the masker, including the fundamental frequency (F0), temporal fine structure (TFS) and the intelligibility of the masking voice. Differences in F0 and TFS between the target speech and the masker can improve speech intelligibility by facilitating the separation of speech streams (Francart et al., 2011). Zekveld et al. (2014) confirmed these effects. They showed that same-sex speakers cause more spectral overlap, making separation harder, while mixed-sex speakers reduce overlap, facilitating speech perception. They also suggested that spatial separation is especially beneficial for same-sex speakers. Brungart (2001, 2005) confirmed that speech intelligibility is better with mixed-sex speakers and worst when the same speaker is used as both target and masker. Based on the literature, lower spectral overlap is expected to result in better AAD performance, thus resulting in better AAD performance in mixed sex conditions compared to same sex conditions. However, **we hypothesized that AAD performance will be better in same-sex conditions, as these require more listening effort and therefore might result in higher AAD scores.** This is based on studies that showed that slightly more difficult scenarios can yield higher AAD scores (Das et al., 2016, 2018).

The main goal of this study was to investigate which factors influence AAD performance. We focused on the following research questions:

‐ What is the effect of level of background noise (4dB SNR compared to no noise) on AAD performance?
‐ What is the effect of the speaker’s sex (target and masker) on AAD performance?
‐ What is the effect of listener’s sex on AAD performance?
‐ What is the difference in AAD performance for same compared to mixed-sex conditions?

## 2. Methods

### 2.1 Participants

Forty-two subjects participated, after providing informed consent, of which three were excluded from the analysis. The data of subjects 20 and 25 were excluded due to a technical issue (no noise conditions were played) which made the data unreliable. The data of subject 33 was removed from the analysis because this subject did not understand the task correctly. The subject listened to the wrong speakers which made the ground truth of attention invalid. We removed a few trials from some of the remaining subjects, because of either wrong attention of the subject (3 trials) which we were able to track down in the behavioural questions after the conditions, or a technical issue (in 8 trials) with the triggers of the EEG.

As a result, thirty-nine normal-hearing (≤ 25dB HL), (18;08 – 55;06 years, with median age 21;07 years) adults were included in this study, of which 23 were female and 16 male. Hearing thresholds were confirmed with pure tone audiometry with a threshold *≤* 25dB HL at all octave frequencies from 125 Hz to 8000 Hz. The ear canal and tympanic membrane were checked with otoscopy. In addition, speech intelligibility was measured with a speech-in-noise test using MATRIX sentences (Luts et al., 2014). The native language of the participants was Dutch (Flemish). Subjects with neurological disorders (cognitive or attentional), such as ASS, AD(H)D, dyslexia, dyscalculia, epilepsy were excluded from the research. At the start of the study, participants completed a questionnaire addressing the inclusion and exclusion criteria, such as age and relevant disorders. The study was conducted under approval of the Medical Ethics Committee UZ Leuven/Research (KU Leuven) with reference S57102.

### 2.2 Apparatus

The measurements took place in a soundproof, electromagnetically shielded booth. Participants were seated in a comfortable chair in front of a large screen with a projector. Behind the screen, 3 loudspeakers (Genelec type 8020A) presented the stimuli and were positioned at -45°, 0°, and 45°. A fourth loudspeaker (Fostex type 6031B) was located above the participant (see Figure 1).

**Figure 1.**
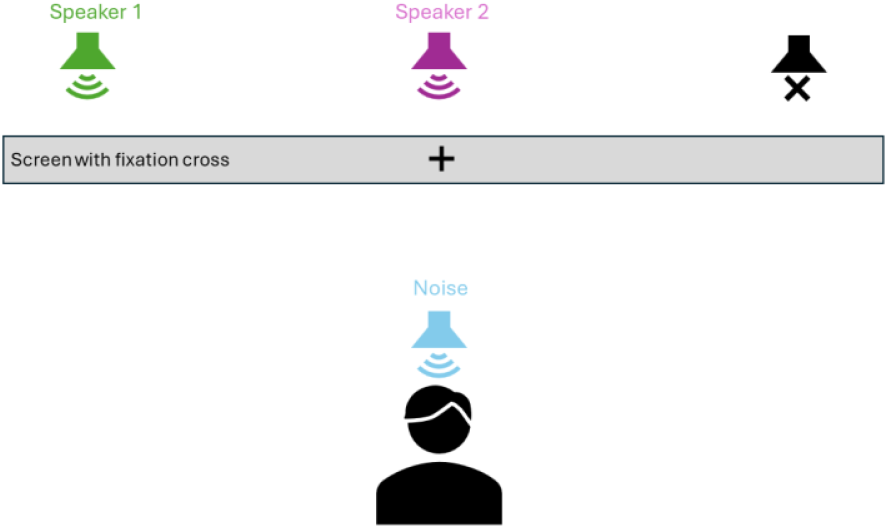
Visual representation of the set-up

Stimuli were presented using APEX 4.1.2 (Francart et al., 2008) and an RME Multiface II sound card (Haimhausen, Germany). The sound levels were calibrated using a 2 cm^3^ coupler of the artificial ear (Brüel&Kjaer 4152, Denmark). Communication could be maintained with the participant via a microphone in the booth.

Brain activity was recorded with 64-channel Biosemi ActiveTwo EEG with a sampling rate of 8192 Hz. Participants were instructed to gaze at a black screen with a white fixation cross during all the conditions. Eye gaze was measured via electro-oculography (EOG) using 4 Biosemi Flat-Type Active electrodes.

Before the measurement, participants were instructed to move as little as possible to avoid muscle artifacts in the EEG signals.

### 2.3 Stimuli

Throughout the experiment, two simultaneous speakers were presented at 60 dBA. The presented stimuli were selected podcasts from the ‘Universiteit van Vlaanderen’ series (*Podcasts Universiteit van Vlaanderen*, n.d.). There was always one speaker per podcast. The stimuli were pre-processed in MATLAB. The intro and outro were cut, and silences were cut to a maximum of 500 milliseconds. Additionally, the podcasts were normalized to the same RMS level. The fundamental frequency (F0) of the male speakers was lower than 193 Hz and of the female speakers always higher than 193 Hz. The noise which was presented by a speaker above the participants was a long-term average speech spectrum (LTASS) weighted noise, where the LTASS was computed from the selected podcasts.

### 2.4 Conditions

The 2 speakers were consecutively presented from 3 different combinations of locations: -45°/0° (left, front), 0°/45° (front, right) or -45°/45° (left, right). All participants ran through the same 10 conditions.

The total experiment time was 180 minutes, of which 65 minutes were the EEG measurement consisting of 10 conditions. Five-minute breaks were obligatory after condition 3 and 6. In addition, after each condition participants were asked if they needed a break. Each condition lasted 6min30s. The first tryout condition and 8 experimental conditions contained 5 switches of the locus of attention, each happening every minute. The participants attended to one podcast the entire condition, but this podcast switched position every minute. These locus switches are shown in Figure 2. There are 6 different types of locus of attention combinations, L/F, F/L, L/R, R/L; R/F, F/R, with L= left (-45°), F= front (0°) R= right (45°).The tenth condition was the Reference condition, in which only 1 switch of the locus of attention was present halfway that condition.

**Figure 2.**
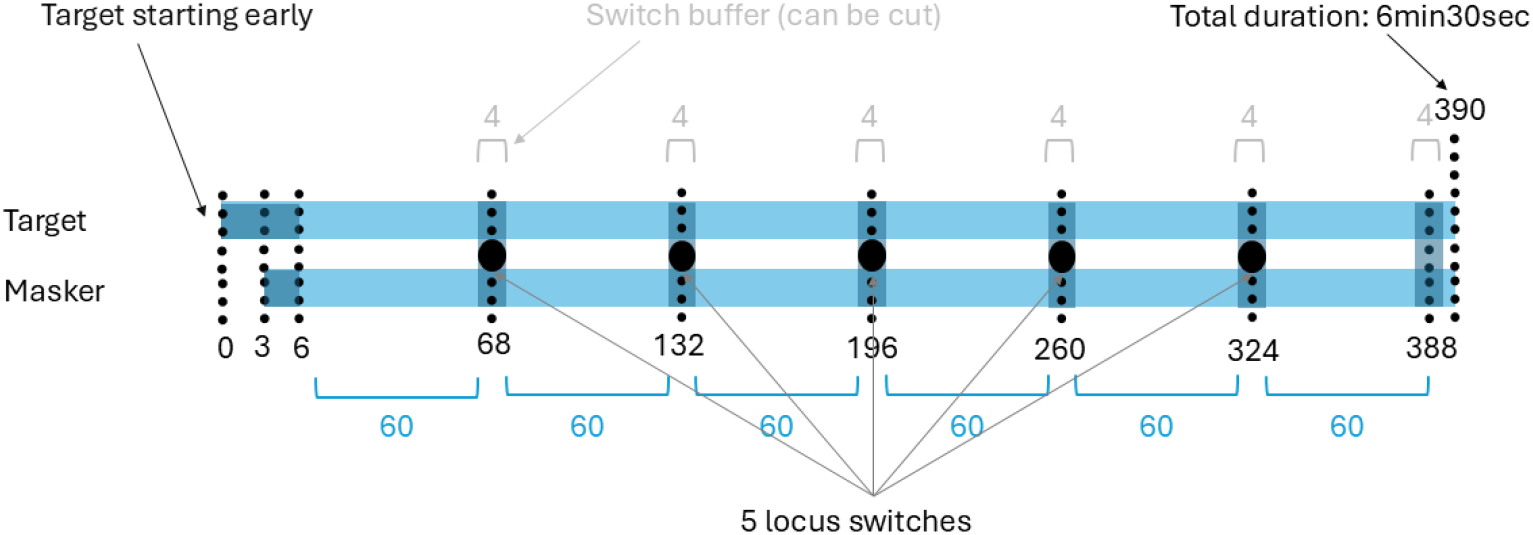
The timeline of a representative condition. All values are expressed in seconds. The dots indicate that every minute the locus of attention changes. The Target stimulus starts a few seconds early to avoid confusion as to which stimulus the participant should listen to. Each participant listened to 10 conditions of 6 min30s, so 65 min in total of speech during the entire EEG session.

In total, there were 51 podcasts by female speakers and 59 podcasts by male speakers. For each subject, 8 podcasts with female voices and 8 podcasts with male voices were drawn randomly from the pool of 51 and 59 podcasts without replacement. In addition, the attended/unattended locus of the 2 active speakers was never the same twice in a row (i.e. there was always a real locus switch). Moreover, the same noise level, sex of the Target speaker, and sex of the Masker speaker never appeared three times in a row. Subjects were instructed to attend to 1 podcast/speaker (i.e. the Target speaker) and ignore the other, unattended speaker (i.e. the Masker speaker). In every condition, participants were reminded about the instructions of the task via the screen. This instruction showed, before the start of each condition, where the Target speaker would initially be presented (left, front or right loudspeaker/position).

The first condition was always the Tryout condition, which served to get acquainted with the procedure. Then 8 experimental conditions were presented (in a randomized order). Those conditions were divided into 4 conditions with a background noise of +4dB SNR and 4 conditions without background noise (in silence). Four conditions had Same-Sex (Target Male - Masker Male, Target Female - Masker Female) and the other 4 had Mixed-Sex (Target Male - Masker Female, Target Female - Masker Male) configurations. These conditions are represented in Figure 3.

**Figure 3.**
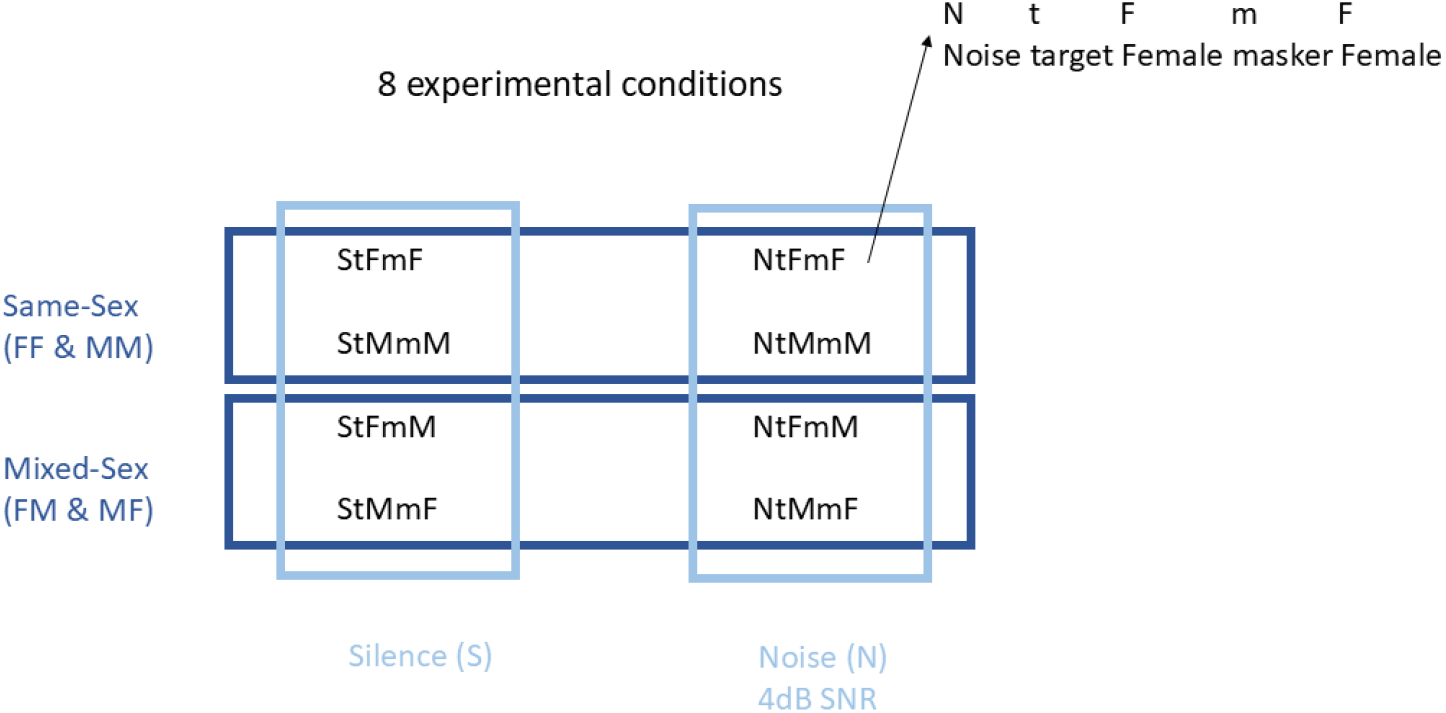
Overview of the 8 experimental conditions. The speaker sexes are organised in order with first the Target speaker and then the Masker speaker.

The last condition was always the reference condition. This was the only condition that differs in the number of switches of the locus of attention. Instead of 1 switch every minute (5 switches in total), there was only 1 switch in the middle of the condition. This condition served as a control condition (standard AAD). The tryout and reference conditions had the same stories but in the opposite order, i.e. the Masker speaker from the tryout condition became the Target speaker in the reference condition and the Target speaker from the tryout condition became the Masker speaker in the reference condition. Only the 8 experimental conditions are included in the analysis in the Results section.

All participants received the same instructions, presented by means of an instructional video at the beginning of the test session, which explained the procedure. After each condition, there were multiple choice questions about the content of the story to keep the participants’ attention and motivation maximum. Furthermore, participants were asked an open question about their experience to find out whether the ground truth of attention was correct. The question was “What did you think of this condition? How did you experience it? Do you have any comments/observations? This can be about anything, e.g. the content of the podcast, listening itself, your fatigue etc.”

### 2.5 Signal processing

#### 2.5.1 Envelope Reconstruction/Processing/analysing EEG data

To investigate the effects of the speaker and subject-related factors, the AAD decoding accuracies were calculated. The EEG data was processed offline in MATLAB (R2016b), after the measurement.

##### Speech stimuli

The speech envelope was extracted from the original speech signal following the method outlined by Biesmans et al. (2017) utilizing a gamma-tone filter bank (using 28 channels spaced by 1 equivalent rectangular bandwidth with center frequencies ranging from 50 to 5000 Hz), applying a power law of 0.6, and subsequently band-pass filtering the envelope in the delta (0,5-4 Hz) and theta (4-8 Hz) frequency bands. The 28 bands were averaged to obtain one envelope signal. Then, the speech envelope was downsampled to 128 Hz to match the frequency of the downsampled EEG signal.

##### EEG processing

The EEG data was collected at a sampling rate of 8192 Hz. The data was processed per condition. The EEG speech envelope was reconstructed using a linear decoder, estimated via ridge regression with the mTRF toolbox, similar to Crosse et al. (2016). This approach accounted for all 64 EEG channels and their time-lagged variants from 0 to 250 ms. Data was removed around the attention switches within each condition, namely 2 seconds before and 2 seconds after the switch (see ‘switch buffer’ in Figure 2). This was done to ensure a correct ground truth of attention. Before the envelope reconstruction, the following signal processing steps were applied to the EEG data: First the data was downsampled from 8192 Hz to 256 Hz using MATLAB’s resample function with an anti-aliasing filter to decrease preprocessing time. Then, the spatio-temporal characteristics of eye-blink artifacts were estimated and removed using a Multichannel Wiener filter (MWF), as described by Somers et al. (2018). Then, a bandpass Chebyshev filter was applied, which retained frequencies between 1 and 9 Hz, as these encompass most of the dominant speech features. After the bandpass filter all channels are rereferenced to the common average reference (the reference is the mean of all the EEG channels), followed by a second downsampling step to 128 Hz.

#### 2.5.2 AAD-decoder

Only the data from the 8 experimental conditions was used in the analysis, i.e. the tryout and reference conditions were not included.

For AAD, we have used a backward linear decoder model to reconstruct the speech envelope from the multi-channel EEG (Crosse et al., 2016). The reconstructed envelope of the attended source from the EEG is correlated with the original stimulus envelopes (Crosse et al., 2016) and the one with the highest correlation is considered the attended one. A decision window length of 30 seconds was chosen, which refers to the duration of the window used to correlate the reconstructed envelope with the original stimuli envelopes.

The decoder was trained with the **6-fold Cross-Validation (CV)** method (See Figure 4). A segment is defined here as a block of EEG of 1 minute. The decoder was trained **subject specifically** on an equal amount of data from each condition (**balanced** across conditions). In each CV fold, the backward models were trained on 5 minutes of data per condition (50 minutes of training data in total) and predict who the attended speaker is in the remaining minute of each condition. This was repeated 6 times, such that the attended speaker was predicted once for each segment in the dataset. These predictions were then aggregated per condition, leading to a single prediction accuracy (%) per condition per subject.

**Figure 4.**
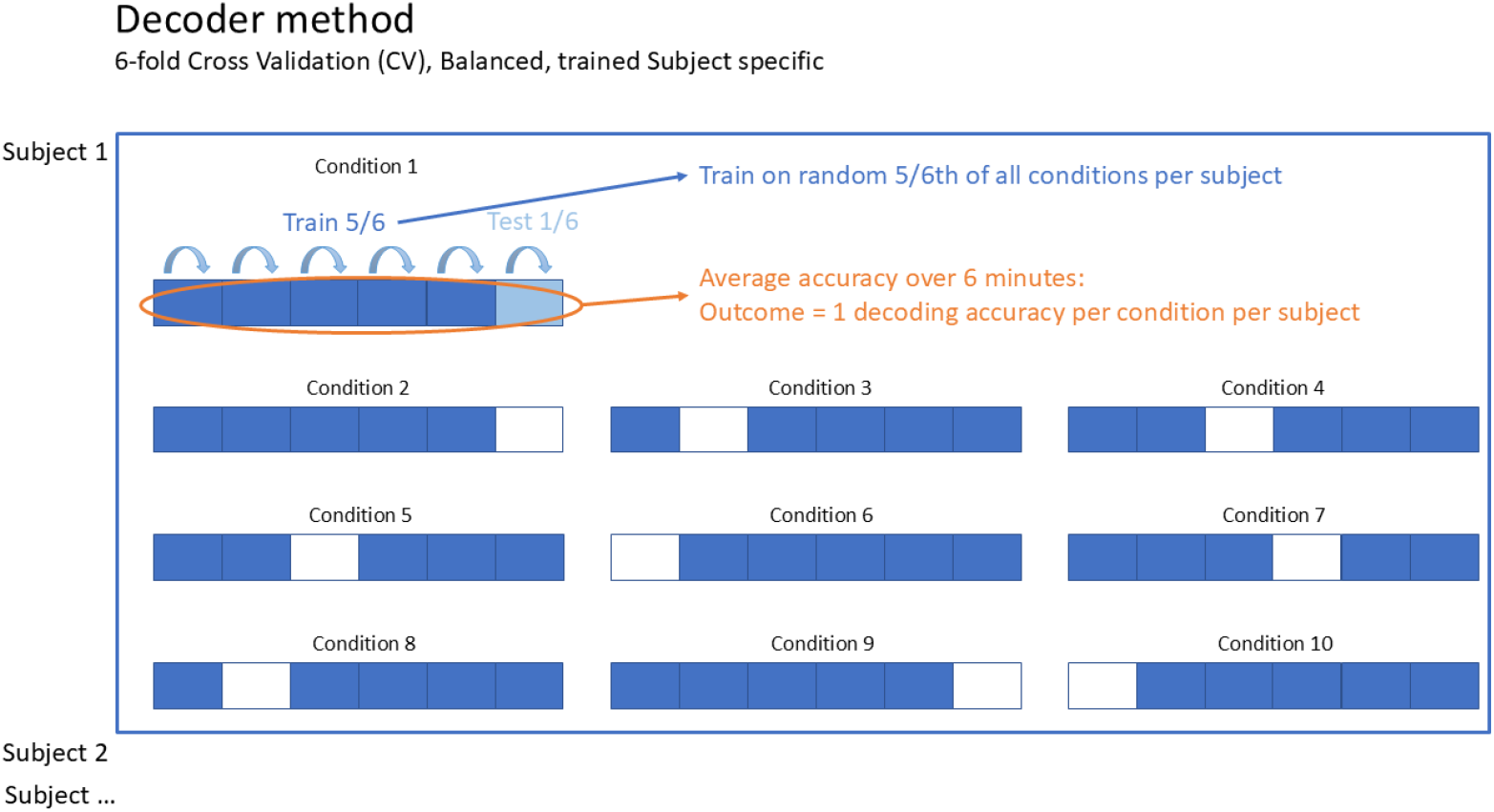
This figure shows an example fold the 6-fold cross-validation (CV) with a balanced design across conditions (in each CV fold, a single 1-minute segment was randomly selected in each condition as test data, while randomly chosen 5/6th of each condition of the rest of the data was used as training data).

### 2.6 Statistical analysis

Statistical analysis was performed using R (version 3.6.3) software. The significance level was set at α=0.05. The chosen statistical method to investigate the sex and noise effects on AAD performance was a Linear Mixed Effects (LME) Model (Field et al., 2012; Galecki & Burzykowski, 2012; Winter, 2013). The LME model had the following general formula:

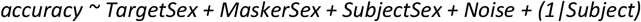

The dependent variable was AAD decoding accuracy (%). The random effect was subject. The fixed effects were Sex of the Masker speaker (M/F), Sex of the Target speaker (M/F), Sex of the Listener (M/F), Same/Mixed-sex and Noise (Noise/Silence), which are factorial predictors, with the letters M representing Male and F representing Female.

The LME models for the interaction effects (see Table 1 b) had the following general formulas:

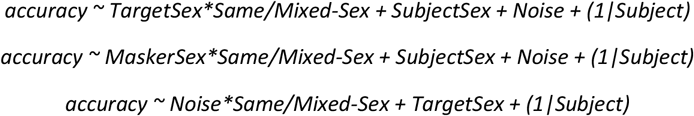

**Table 1.** (a) Overview of the factors with df, t and p values, as well as the 95% - Confidence Intervals, showing a significant p-value for Target and Masker speakers. (b) shows the interaction effects between the factors. There is a significant interaction effect for TargetSex, MaskerSex and Noise with the factor Same/Mixed-Sex.

| Factor | df | t | p | Effect | Predicted mean | Contrast estimate | CI of the difference in predicted means |
| --- | --- | --- | --- | --- | --- | --- | --- |
| Sex Target speaker | 279,138 | 2,091 | 0,0374* | M > F | Male: 0,802<br>Female: 0,773 | -0,0284 | [-0.055 -0.002] |
| Sex Masker speaker | 279,184 | 2,386 | 0,0177* | M > F | Male: 0,804<br>Female: 0,771 | -0,0324 | [-0.059 -0.006] |
| Sex Listener | 39,771 | -1,063 | 0,2943 | / | Male: 0,772<br>Female: 0,803 | 0,0308 | [-0.028 0.089] |
| Level Background Noise | 279,478 | 0,452 | 0,6517 | / | Noise: 0,784<br>Silence: 0,790 | -0,0061 | [-0.033 0.021] |

(b) Interaction effects
| Factors | df | t | p- | Effect | Predicted mean |
| --- | --- | --- | --- | --- | --- |
| Same/Mixed * Target | 278,180 | 2,383 | 0,0178* | In Same-Sex<br>MM > FF | female mixed: 0,791 |
|  |  |  |  |  | male mixed: 0,787 |
|  |  |  |  |  | female same: 0,755 |
|  |  |  |  |  | male same: 0,816 |
| Same/Mixed * Masker | 278,133 | 2,089 | 0,0376* | In Same-Sex<br>MM > FF | female mixed: 0,787 |
|  |  |  |  |  | male mixed: 0,791 |
|  |  |  |  |  | female same: 0,755 |
|  |  |  |  |  | male same: 0,816 |
| Same/Mixed *<br>Noise | 278,490 | 2,865 | 0,0045** | Silence > Noise<br>in Same-Sex | noise mixed: 0,807 |
|  |  |  |  |  | silence mixed: 0,774 |
|  |  |  |  |  | noise same: 0,765 |
|  |  |  |  |  | silence same: 0,811 |

| Effect | Contrast | Contrast estimate | CI of the difference in predicted means |
| --- | --- | --- | --- |
| Same/Mixed Target * | female mixed – male mixed | 0,00401 | [-0.034 0.042] |
|  | female mixed – female same | 0,03538 | [-0.003 0.073] |
|  | female mixed – male same | -0,02551 | [-0.063 0.012] |
|  | male mixed – female same | 0,03138 | [-0.007 0.069] |
|  | male mixed – male same | -0,02952 | [-0.067 0.008] |
|  | female same – male same | -0,06089 | [-0.099 -0.023] |
| Same/Mixed Masker * | female mixed – male mixed | -0.00401 | [-0.042 0.034] |
|  | female mixed – female same | 0.03138 | [-0.007 0.069] |
|  | female mixed- male same | -0.02952 | [-0.067 0.008] |
|  | male mixed – female same | 0.03538 | [-0.003 0.073] |
|  | male mixed – male same | -0.02551 | [-0.063 0.012] |
|  | female same – male same | -0.06089 | [-0.099 -0.023] |
| Same/Mixed * Noise | female mixed – male mixed | -0.00401 | [-0.042 0.034] |
|  | female mixed – female same | 0.03138 | [-0.007 0.069] |
|  | female mixed – male same | -0.02952 | [-0.067 0.008] |
|  | male mixed – female same | 0.03538 | [-0.003 0.073] |
|  | male mixed – male same | -0.02551 | [-0.063 0.012] |
|  | female same – male same | -0.06089 | [-0.099 -0.023] |

The predicted means of the main and interaction effects were calculated with the emmeans() function in R.

## 3. Results

Figure 5 shows the boxplots of the 8 experimental conditions.

**Figure 5.**
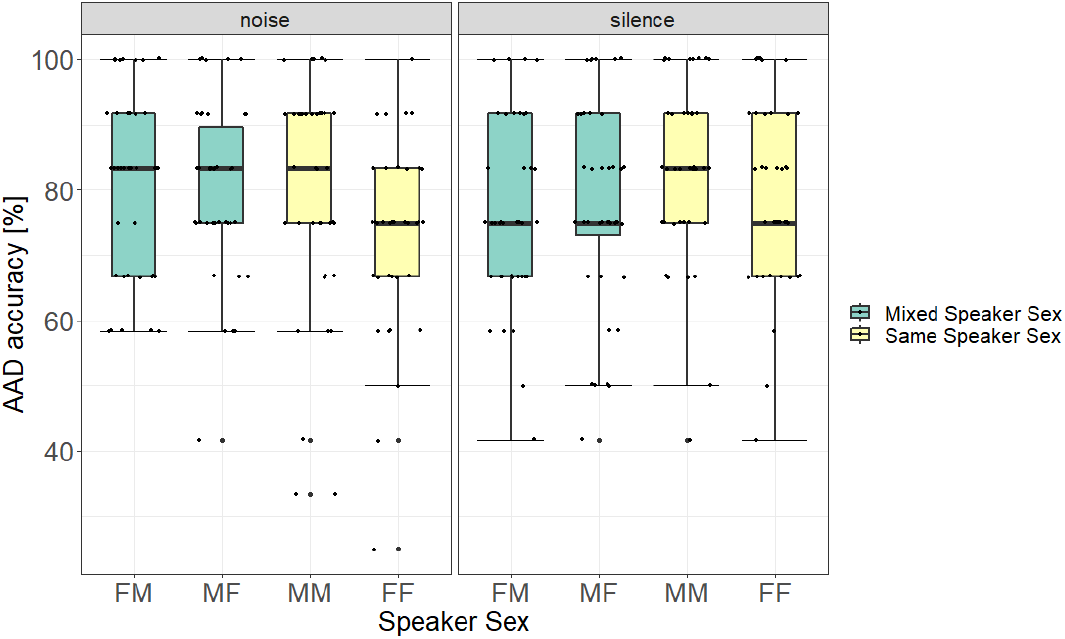
Overview of the 8 experimental conditions, showing the distribution of the conditions in levels of background noise, Same/Mixed-sex and sex of the Target and Masker speaker.

We did not observe a significant main effect of Sex of the Listeners (See Figure 6), Same/Mixed-Sex, nor from the level of Background noise. However, a significant main effect was found for the Sex of the Target Speaker and also of the Sex of the Masker Speaker. The results are presented in Table 1. Additionally, Figure 6 shows the same data grouped by sex of the Target or Masker. It shows that AAD accuracy is highest for Male Target speakers and Male Masker speakers.

**Figure 6.**
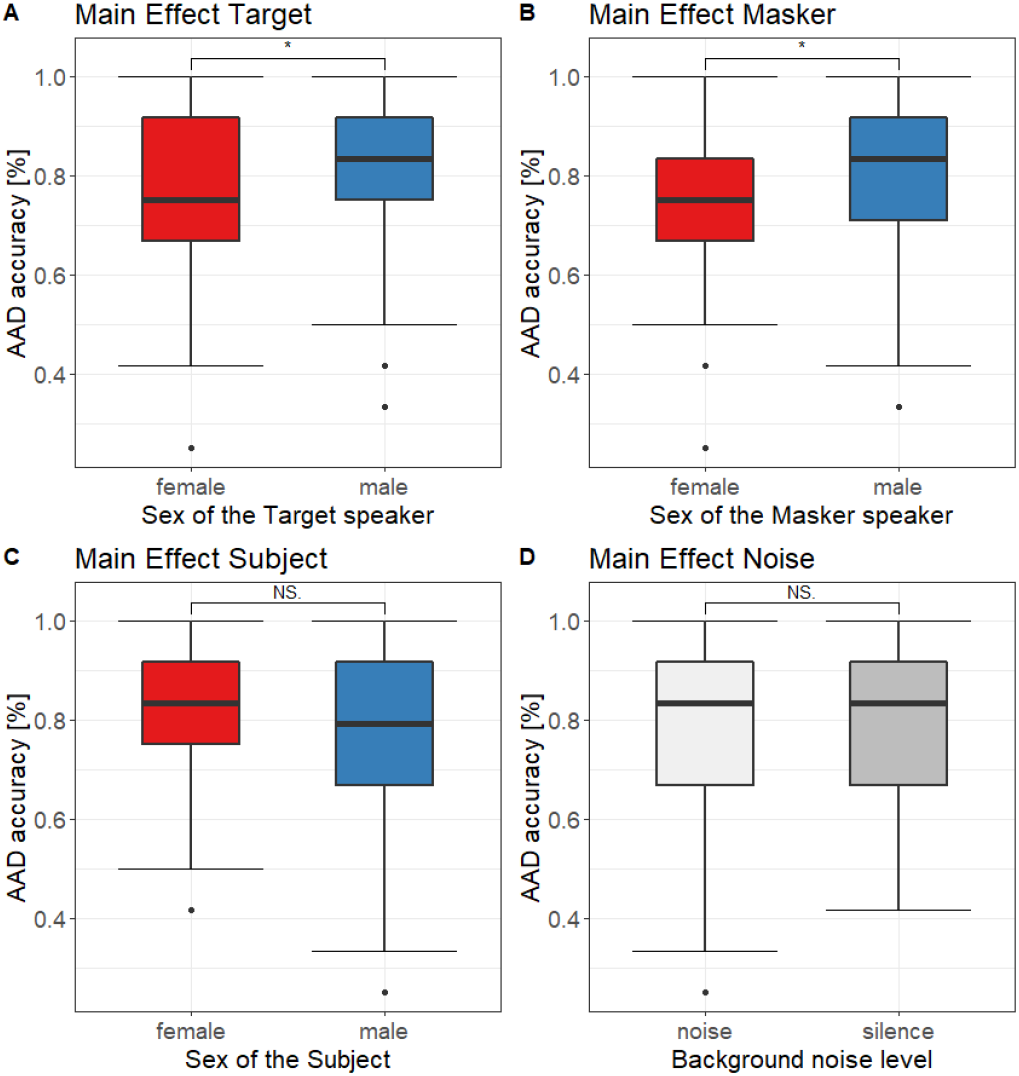
Main effects of factors Target (A), Masker (B), SubjectSex (C) and Noise (D). with female speakers or subjects in red and male speakers or subjects in blue, noise in light grey and silence in dark grey. AAD accuracy is shown as a function of the 4 different factors. A and B show the significant main effect of Target and Masker. AAD accuracy is highest for Male Target speakers and Male Masker speakers. C and D show the non-significant main effects of Sex of the Listener and Noise

We found an additional interaction effect between Same/Mixed-sex and Speaker sex (Target and Masker). In Same-Sex conditions, the conditions with male voices (MM) had higher AAD accuracies than the female conditions (FF). Also in the Same-Sex conditions, there was an interaction effect with the level of background noise. Silence conditions had higher AAD accuracies than noise conditions. In addition, in the Mixed-Sex conditions, noise conditions had higher AAD accuracies than silence conditions. The boxplots showing the interaction effects are shown in Figure 7. The statistical values of the interaction effects are shown in Table 1. Additionally, Table 1 presents the predicted means, contrasts and confidence intervals of the differences in predicted means for the main and interaction effects. These values indicated that the effects were mostly smaller than 5%. We assume that a potential effect smaller than 5% would not have a meaningful practical effect on an AAD system.

**Figure 7.**
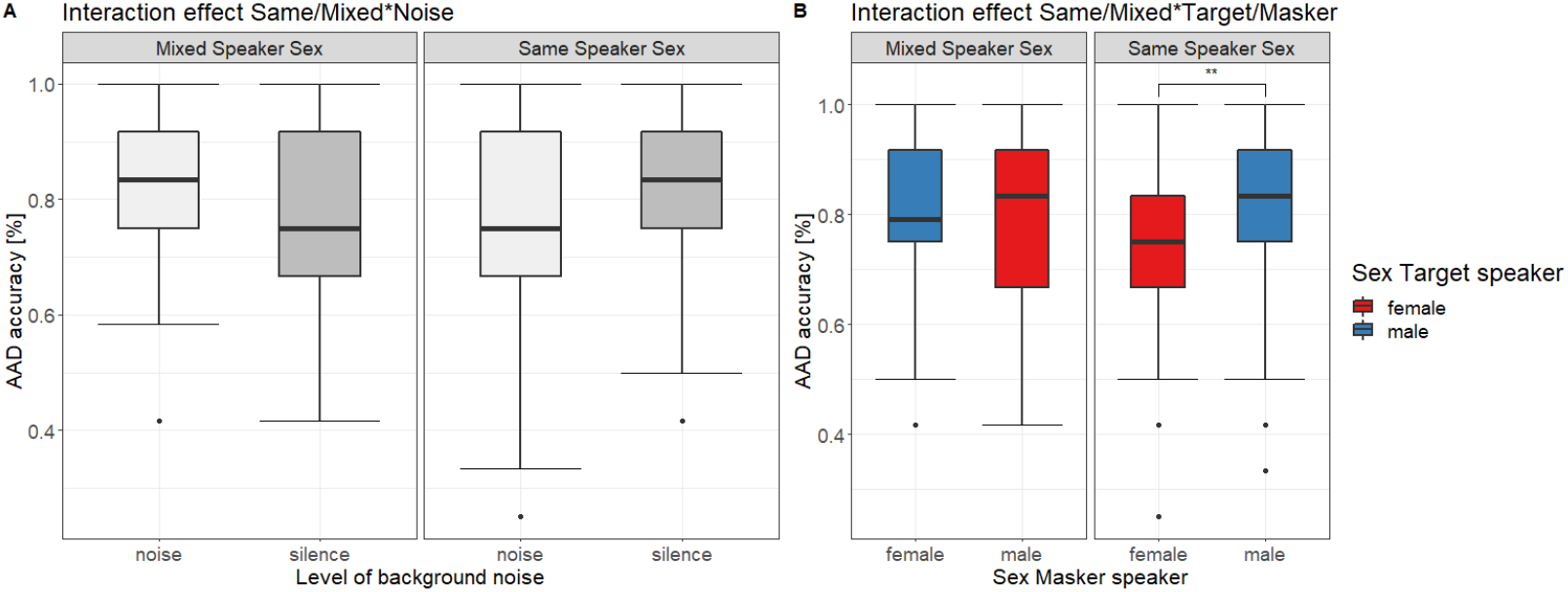
shows the interaction effects of Same/Mixed-Sex with Level of background noise in panel A and with Speaker sex (Target and Masker) in panel B.

## 4. Discussion

Currently, the factors influencing AAD performance remain mostly unknown. The objective of this research was to examine factors that impact AAD performance. We investigated the effect of the Sex of the Speaker, Sex of the Listener, Same versus Mixed-Sex, and background noise level, on AAD performance. In our study, 42 young participants with normal hearing were asked to focus on one speaker (attended) while ignoring another (unattended) during the simultaneous presentation of two podcasts.

### Research question 1 – Background noise level

The first objective of the study was to investigate the effect of background noise levels on AAD performance. Specifically, the question posed was whether AAD performance is better in conditions with a background noise level of +4dB SNR in comparison to conditions without any background noise. In the study conducted by Das et al. (2018) it was observed that in conditions with -1.1 dB SNR, AAD performance was better compared to both conditions in silence and conditions with higher noise levels (-4.1 and -7.1 dB SNR), hypothesizing that adding a small amount of background noise increases listening effort and subsequently improves AAD performance. Therefore, we hypothesized in our study that AAD performance would be better in conditions with a background noise of +4 dB SNR compared to conditions without background noise. Remarkably, our results showed no significant main effect of background noise. The main differences between our protocol and the experimental set-up of Das et al. (2018) were the noise levels (+4dB versus -1,1; -4,1; -7,1; +20dB SNR, respectively), the type of noise used (stationary noise (LTASS) versus babble noise), the method of presenting the stimuli (loudspeakers versus insert phones), the sex of the speakers (all sex combinations versus only MM), the spatial separation (-45°, 0°, 45° versus 10°, 60°, 180°). Moreover, we had a positive noise level (+4dB SNR) which could indicate that the condition was not difficult enough to find a difference with the condition without background noise. The inclusion of background noise in future fitting protocols is crucial, as real-life situations typically involve some level of background noise. It is encouraging that the addition of a limited amount of background noise (in our case 4dB) does not have a large effect on AAD performance, since we aim for AAD to perform well in realistic scenarios. We hypothesize that there might be an interaction between listening effort, noise level and AAD performance.

### Research question 2 – Sex of the speaker

A second research aim was to assess if AAD performance is better with a male or a female speaker. Currently, there is no observed effect of speaker sex on AAD performance in the literature. Within the literature, certain studies have concentrated on the effects of speaker sex on neural auditory processing (Ahadi et al., 2014; Durrant et al., 1990; Jerger & Johnson, 1988; Krizman et al., 2012; Van Canneyt et al., 2021; Weston et al., 2015; Zakaria et al., 2019), while others focused on the effect on speech intelligibility (Gruber & Gaebelein, 1979; Kiliç & Ögüt, 2004; Kwon, 2010). There is no consensus in the literature regarding the effects of speaker sex on neural auditory processing or behavioural measures. We hypothesized to find better AAD performance for male speakers (Target & Masker speaker). Our findings demonstrate that the sex of both the Target and Masker speaker significantly affected AAD decoding accuracy. AAD accuracy is highest for Male Target speakers and Male Masker speakers. This could imply that attending a male voice is easier, but that does not necessarily imply that it is harder to ignore a male voice. These results suggest that a male-male (MM) condition should render the highest AAD accuracies out of all the Same/Mixed-Sex conditions. This was proven in research question 4, showing a significant interaction effect between Same/Mixed-Sex and Speaker Sex.

The effect of Speaker Sex identified in our study is not desirable for the application of neuro-steered hearing aids. It is essential for AAD to perform equally well across all sexes in realistic circumstances.

The confidence intervals of the difference in predicted means indicated that the effect was small and therefore clinically negligible.

### Research question 3 – Sex of the listener

A third research question was determining if AAD performance is better for a male or female listener. In 1-speaker scenarios we found no evidence in the literature about speech intelligibility differences for male and female listeners (Ellis et al., 1996; Hazan & Markham, 2004; Yoho et al., 2019). For ABR’s there is a sex effect. Important to note is that in these papers the performance was measured via speech intelligibility tests instead of brain activity measures. There are gender-specific normative data for ABR in clinical applications (Hall, 1992). Also other studies confirm that neural encoding of the acoustic cues of speech stimuli is different at a subcortical level between male and female **subjects**. (Ahadi et al., 2014; Durrant et al., 1990; Jerger & Johnson, 1988; Krizman et al., 2012; Zakaria et al., 2019). We hypothesized there would be no difference in decoding performance for a male or female listener. In line with our hypothesis, we did not observe a significant main effect of the sex of the listener. Again, the confidence intervals of the difference in predicted means showed that the effect of factor listener sex was small and therefore not clinically meaningful.

### Research question 4 – Same versus Mixed-Sex

A fourth and final research aim explored whether AAD performance is better in Mixed-Sex conditions compared to Same-Sex scenarios. Previous research has demonstrated that in conditions with Mixed-Sex, speech intelligibility was better. It is easier to disentangle the speakers when there is less spectral overlap (Brungart, 2001; Brungart et al., 2005; Zekveld et al., 2014). A study of Francart et al. (2011) indicated that F0 plays a key role in stream segregation, and differences in F0 help in separating speakers, but similar F0s can cause reduced speech recognition. This implies that differentiating between speakers is easier in Mixed-Sex conditions and thus facilitates the tracking of the attended speaker. In addition, there was also informational masking, which is when listeners fail to identify the signal amid the confusion of masking sounds. It is the interference in understanding a speech signal that cannot be accounted for simply by the spectro-temporal overlap of competing sounds. If the cues of the speech signal and Masker are too similar, this can confuse the listener, resulting in lower speech recognition performance (Francart et al., 2011).

However, we hypothesized that AAD performance would be better for Same-Sex conditions since these conditions require more listening effort and therefore might result in higher AAD scores. This hypothesis was based on studies that showed that slightly more difficult scenarios yield higher AAD scores (Das et al., 2016, 2018)

No significant main effect was found of Same compared to Mixed-Sex conditions on AAD performance. However, we did find an additional **interaction effect** between Same/Mixed-Sex and Speaker Sex (Target and Masker). In Same-Sex conditions, the conditions with male voices (MM) had higher AAD accuracies than the female conditions (FF). This is in line with the results that conditions with male speakers have higher AAD performance. The differences in predicted means’ confidence intervals suggested that the effects were minimal and therefore not clinically meaningful.

### Sex summary

From a clinical perspective, the presence of an effect of speaker or listener sex is undesirable. The AAD decoder should perform well in all circumstances.

The lack of significance in the main effect level of background noise may be attributed to an interaction effect with the other factors. In the Same-Sex conditions, there was an interaction effect with the level of background noise. Silence conditions had higher AAD accuracies than noise conditions.

Consequently, AAD performance is better in conditions in silence instead of adding noise on top of this already challenging condition. Note that the confidence intervals of the differences in predicted means showed that the effects were small and therefore clinically negligible.

It is important to emphasize that attention is likely not only influenced by Speaker’s Sex. Numerous other factors related to the speakers and podcasts could have been of influence, such as the topic of the podcasts, participant’s motivation, levels of fatigue, familiarity with the topic, educational background (particularly considering the highly scientific nature of the topics), and cognition.

### Limitations of the study

A factor that could potentially contribute to higher AAD scores is the content of the podcasts. If subjects lacked interest in the topics, it might have resulted in lower scores. To address this in future research, participants could be questioned beforehand about their preferences regarding podcast topics, to better engage their attention. Additionally, participants could be asked whether they had previously listened to the podcast.

To address participant’s reported fatigue during measurements, future studies may consider incorporating tiredness scales to assess fatigue levels. This approach would allow for an investigation into potential correlations between decreased AAD scores and fatigue.

Future research might explore other populations than our homogenous group of young, normal-hearing subjects. Including individuals of various ages and with different degrees of hearing loss can shed light on the impact of these factors on AAD performance. These diverse groups are particularly relevant as they represent the eventual users of neuro-steered aids. Decoders might need to be trained/customized specifically per subject.

Future research may also explore factors that could potentially influence AAD performance, such as the distinction between dialogues and monologues, as well as passive versus active listening or voice familiarity.

## 5. Conclusion

The objective of this research was to examine which factors impact AAD performance, because the AAD decoder should perform well in all circumstances. This study investigated the effect of several factors, such as the sex of the speaker, sex of the listener, Same versus Mixed-Sex speaker pairs, and background noise level, on AAD performance. Notably, we found that the sex of the listener, background noise level, and Same versus Mixed-Sex conditions did not meaningfully impact AAD performance. Nonetheless, our research did indicate that AAD performance is higher for Male Target and Male Masker speakers. While no substantial effects were found on the factors investigated in this study, it remains a good idea to include diverse and realistic training scenarios, to avoid potential other factors influencing the results. Further research is needed to better understand which factors influence AAD performance. These factors might be related to the subject and could include age, degree of hearing loss, voice familiarity, topic familiarity, passive versus active participation, and listening effort.

## Author Statement

Tom Francart, Alexander Bertrand, Debora Fieberg and I.V.d.R. designed the experiment (protocol and practical set-up). Nicolas Heintz and Iustina Rotaru contributed to the analysis of the EEG data and applied the AAD decoders. All authors contributed to discussing the results and implications of the research. I.V.d.R. performed the experiments, did the statistical analysis, and wrote the paper. All co-authors reviewed the paper.

## Acknowledgments

The authors would like to thank Erin Willems and Anouck Jaspers for their help in data acquisition. We would also like to thank all the subjects for their participation in the study.

Declaration of generative AI and AI-assisted technologies in the writing process

During the preparation of this work the author used ChatGPT to rephrase sentences in academically correct English. After using this tool, the author reviewed and edited the content as needed and takes full responsibility for the content of the published article.

## Bibliography

Ahadi, M., Pourbakht, A., Jafari, A. H., Shirjian, Z., & Jafarpisheh, A. S. (2014). Gender disparity in subcortical encoding of binaurally presented speech stimuli: An auditory evoked potentials study. Auris Nasus Larynx, 41(3), 239–243. 10.1016/j.anl.2013.10.010

Aroudi, A., Mirkovic, B., De Vos, M., & Doclo, S. (2019). Impact of Different Acoustic Components on EEG-Based Auditory Attention Decoding in Noisy and Reverberant Conditions. IEEE Transactions on Neural Systems and Rehabilitation Engineering, 27(4), 652–663. 10.1109/TNSRE.2019.2903404

Biesmans, W., Das, N., Francart, T., & Bertrand, A. (2017). Auditory-inspired speech envelope extraction methods for improved EEG-based auditory attention detection in a cocktail party scenario. IEEE Transactions on Neural Systems and Rehabilitation Engineering, 25(5), 402–412. 10.1109/TNSRE.2016.2571900

Biesmans, W., Vanthornhout, J., Wouters, J., Moonen, M., Francart, T., & Bertrand, A. (2015). Comparison of speech envelope extraction methods for EEG-based auditory attention detection in a cocktail party scenario. Proceedings of the Annual International Conference of the IEEE Engineering in Medicine and Biology Society, EMBS, 2015-Novem, 5155–5158. 10.1109/EMBC.2015.7319552

Brungart, D. S. (2001). Informational and energetic masking effects in the perception of two simultaneous talkers. The Journal of the Acoustical Society of America, 109(3), 1101–1109. 10.1121/1.1345696

Brungart, D. S., Simpson, B. D., & Freyman, R. L. (2005). Precedence-based speech segregation in a virtual auditory environment. The Journal of the Acoustical Society of America, 118(5), 3241–3251. 10.1121/1.2082557

Coates, J. (2004). Women, men and language: a sociolinguistic account of gender differences in language (3rd ed.). Harlow: Pearson.

Cohen, J. (1988). Statistical power analysis for the behavioral sciences (Second edi). Lawrence Erlbaum Associates. https://books.google.be/books?id=2v9zDAsLvA0C&pg=PP1&redir_esc=y#v=onepage&q&f=false

Cohen, J. (1992). Quantitative methods in psychology. 112(1), 155–159. 10.1037/0033-2909.112.1.155

Crosse, M. J., Di Liberto, G. M., Bednar, A., & Lalor, E. C. (2016). The multivariate temporal response function (mTRF) toolbox: A MATLAB toolbox for relating neural signals to continuous stimuli. Frontiers in Human Neuroscience, 10(NOV2016), 1–14. 10.3389/fnhum.2016.00604

Das, N., Bertrand, A., & Francart, T. (2018). EEG-based auditory attention detection: Boundary conditions for background noise and speaker positions. Journal of Neural Engineering, 15(6). 10.1088/1741-2552/aae0a6

Das, N., Biesmans, W., Bertrand, A., & Francart, T. (2016). the Effect of Head - Related Filtering and Ear - Specific Decoding Bias. Journal of Neural Engineering, 32(0). 10.1088/1741-2560/13/5/056014

Das, N., Van Eyndhoven, S., Francart, T., & Bertrand, A. (2017). EEG-based attention-driven speech enhancement for noisy speech mixtures using N-fold multi-channel wiener filters. 25th European Signal Processing Conference, EUSIPCO 2017, 2017-Janua, 1160–1164. 10.23919/EUSIPCO.2017.8081390

Durrant, J. D., Sabo, D. L., & Hyre, R. J. (1990). Gender, head size, and abrs examined in large clinical sample. In Ear and Hearing (Vol. 11, Issue 3, pp. 210–214). 10.1097/00003446-199006000-00008

Ellis, L., Fucci, D., Reynolds, L., & Benjamin, B. (1996). Effects of gender on listeners’ judgments of speech intelligibility. Perceptual and Motor Skills, 85(3 PART 1), 771–775. 10.2466/pms.1996.83.3.771

Ericson, M. A., Brungart, D. S., & Simpson, B. D. (2004). Factors that influence intelligibility in multitalker speech displays. International Journal of Aviation Psychology, 14(3), 313–334. 10.1207/s15327108ijap1403_6

Field, A., Miles, J., & Field, Z. (2012). Discovering statistics using R. Sage.

Francart, T., van Wieringen, A., & Wouters, J. (2008). APEX 3: a multi-purpose test platform for auditory psychophysical experiments. Journal of Neuroscience Methods, 172(2), 283–293. 10.1016/j.jneumeth.2008.04.020

Francart, T., van Wieringen, A., & Wouters, J. (2011). Comparison of fluctuating maskers for speech recognition tests. International Journal of Audiology, 50(1), 2–13. 10.3109/14992027.2010.505582

Galecki, A., & Burzykowski, T. (2012). Linear Mixed-effects Models Using R. In Springer (Vol. 8, Issue 1). Springer Science+Business Media New York 2013. 10.1007/978-1-4614-3900-4

Geirnaert, S., Francart, T., & Bertrand, A. (2019). A new metric to evaluate auditory attention detection performance based on a Markov chain. European Signal Processing Conference, 2019-Septe, 1–5. 10.23919/EUSIPCO.2019.8903146

Geirnaert, S., Francart, T., & Bertrand, A. (2020). An Interpretable Performance Metric for Auditory Attention Decoding Algorithms in a Context of Neuro-Steered Gain Control. IEEE Transactions on Neural Systems and Rehabilitation Engineering, 28(1), 307–317. 10.1109/TNSRE.2019.2952724

Gruber, K. J., & Gaebelein, J. (1979). Sex differences in listening comprehension. Sex Roles, 5(3), 299–310. 10.1007/BF00287397

Hall, J. W. (1992). Handbook of Auditory Evoked Responses. Allyn and Bacon.

Han, C., O’Sullivan, J., Luo, Y., Herrero, J., Mehta, A. D., & Mesgarani, N. (2019). Speaker-independent auditory attention decoding without access to clean speech sources. Science Advances, 5(5), 1–11. 10.1126/sciadv.aav6134

Hazan, V., & Markham, D. (2004). Acoustic-phonetic correlates of talker intelligibility for adults and children. The Journal of the Acoustical Society of America, 116(5), 3108–3118. 10.1121/1.1806826

Hillenbrand, J., Getty, L. A., Clark, M. J., & Wheeler, K. (1995). Acoustic characteristics of American English vowels. Journal of the Acoustical Society of America, 97(5), 3099–3111. 10.1121/1.411872

Hillenbrand, J. M., & Michael, J. C. (2009). The role of f0 and formant frequencies in distinguishing the voices of men and women. Attention, Perception, and Psychophysics, 71(7), 1439–1459. 10.3758/APP

Holtze, B., Jaeger, M., Debener, S., Adiloğlu, K., & Mirkovic, B. (2021). Are They Calling My Name? Attention Capture Is Reflected in the Neural Tracking of Attended and Ignored Speech. Frontiers in Neuroscience, 15(March), 1–15. 10.3389/fnins.2021.643705

Jerger, J., & Johnson, K. (1988). Interactions of age, gender, and sensorineural hearing loss on ABR latency. In Ear and Hearing (Vol. 9, Issue 4, pp. 168–176). 10.1097/00003446-198808000-00002

Kiliç, M. A., & Ögüt, F. (2004). The effect of the speaker gender on speech intelligibility in normal-hearing subjects with simulated high frequency hearing loss. Rev. Laryngol. Otol. Rhinol/, June, 35–38.

Krizman, J., Skoe, E., & Kraus, N. (2012). Sex differences in auditory subcortical function. Clinical Neurophysiology, 123(3), 590–597. 10.1016/j.clinph.2011.07.037

Kwon, H. B. (2010). Gender difference in speech intelligibility using speech intelligibility tests and acoustic analyses. Journal of Advanced Prosthodontics, 2(3), 71–76. 10.4047/jap.2010.2.3.71

Leung, Y., Oates, J., & Chan, S. P. (2018). Voice, articulation, and prosody contribute to listener perceptions of speaker gender: A systematic review and meta-analysis. Journal of Speech, Language, and Hearing Research, 61(2), 266–297. 10.1044/2017_JSLHR-S-17-0067

Luts, H., Jansen, S., Dreschler, W., & Wouters, J. (2014). Development and normative data for the Flemish/Dutch Matrix test. December, 1–6.

Mirkovic, B., Bleichner, M. G., De Vos, M., & Debener, S. (2016). Target speaker detection with concealed EEG around the ear. Frontiers in Neuroscience, 10(JUL), 1–12. 10.3389/fnins.2016.00349

O’Sullivan, J. A., Power, A. J., Mesgarani, N., Rajaram, S., Foxe, J. J., Shinn-Cunningham, B. G., Slaney, M., Shamma, S. A., & Lalor, E. C. (2015). Attentional Selection in a Cocktail Party Environment Can Be Decoded from Single-Trial EEG. Cerebral Cortex, 25(7), 1697–1706. 10.1093/cercor/bht355

Pernet, C. R., & Belin, P. (2012). The role of pitch and timbre in voice gender categorization. Frontiers in Psychology, 3(FEB), 1–11. 10.3389/fpsyg.2012.00023 Podcasts Universiteit van Vlaanderen. (n.d.). https://www.universiteitvanvlaanderen.be/podcast

Rotaru, I., Geirnaert, S., Heintz, N., Van de Ryck, I., Bertrand, A., & Francart, T. (2024). What are we really decoding? Unveiling biases in EEG-based decoding of the spatial focus of auditory attention. Journal of Neural Engineering, 21(1). 10.1088/1741-2552/ad2214

Somers, B., Francart, T., & Bertrand, A. (2018). A generic EEG artifact removal algorithm based on the multi-channel Wiener filter. Journal of Neural Engineering, 15(3), 0–12. 10.1088/1741-2552/aaac92

Tanveer, M. A., Skoglund, M. A., Bernhardsson, B., & Alickovic, E. (2024). Deep learning-based auditory attention decoding in listeners with hearing impairment. Journal of Neural Engineering, 21(3). 10.1088/1741-2552/ad49d7

Titze, I. R. (1989). Physiologic And Acoustic Differences Between Male And Female Voices. Journal of the Acoustical Society of America, 85(4), 1699–1707. 10.1121/1.397959

Titze, I. R., & Martin, D. W. (1998). Principles of Voice Production . The Journal of the Acoustical Society of America, 104(3), 1148–1148. 10.1121/1.424266

Van Canneyt, J., Wouters, J., & Francart, T. (2021). Neural tracking of the fundamental frequency of the voice: The effect of voice characteristics. European Journal of Neuroscience, 53(11), 3640–3653. 10.1111/ejn.15229

Van Eyndhoven, S., Francart, T., & Bertrand, A. (2017). EEG-Informed Attended Speaker Extraction from Recorded Speech Mixtures with Application in Neuro-Steered Hearing Prostheses. IEEE Transactions on Biomedical Engineering, 64(5), 1045–1056. 10.1109/TBME.2016.2587382

Wang, L., Wu, E. X., & Chen, F. (2021). EEG-based auditory attention decoding using speech-level-based segmented computational models. Journal of Neural Engineering, 18(4). 10.1088/1741-2552/abfeba

Weston, P. S. J., Hunter, M. D., Sokhi, D. S., Wilkinson, I. D., & Woodruff, P. W. R. (2015). Discrimination of voice gender in the human auditory cortex. NeuroImage, 105, 208–214. 10.1016/j.neuroimage.2014.10.056

Winter, B. (2013). Linear models and linear mixed effects models in R with linguistic applications. ArXiv, 1–42. http://arxiv.org/pdf/1308.5499.pdf

Wong, D. D. E., Fuglsang, S. A., Hjortkjær, J., Ceolini, E., Slaney, M., & de Cheveigné, A. (2018). A comparison of regularization methods in forward and backward models for auditory attention decoding. Frontiers in Neuroscience, 12(AUG), 1–16. 10.3389/fnins.2018.00531

Xu, Z., Bai, Y., Zhao, R., Hu, H., Ni, G., & Ming, D. (2022). Decoding selective auditory attention with EEG using a transformer model. Methods, 204(April 2022), 410–417. 10.1016/j.ymeth.2022.04.009

Xu, Z., Bai, Y., Zhao, R., Zheng, Q., Ni, G., & Ming, D. (2022). Auditory attention decoding from EEG-based Mandarin speech envelope reconstruction. Hearing Research, 422, 108552. 10.1016/j.heares.2022.108552

Yoho, S. E., Borrie, S. A., Barrett, T. S., & Whittaker, D. B. (2019). Are there sex effects for speech intelligibility in American English? Examining the influence of talker, listener, and methodology. Attention, Perception, and Psychophysics, 81(2), 558–570. 10.3758/s13414-018-1635-3

Zakaria, M. N., Wahab, N. A. A., Maamor, N., Jalaei, B., & Dzulkarnain, A. A. A. (2019). Auditory brainstem response (ABR) findings in males and females with comparable head sizes at supra-threshold and threshold levels. Neurology Psychiatry and Brain Research, 32(May 2018), 4–7. 10.1016/j.npbr.2019.03.001

Zekveld, A. A., Rudner, M., Kramer, S. E., Lyzenga, J., & Rönnberg, J. (2014). Cognitive processing load during listening is reduced more by decreasing voice similarity than by increasing spatial separation between target and masker speech. Frontiers in Neuroscience, 8(8 APR), 1–11. 10.3389/fnins.2014.00088

